# Context-Aware Deep Learning Enables High-Efficacy Localization of High Concentration Microbubbles for Super-Resolution Ultrasound Localization Microscopy

**DOI:** 10.1101/2023.04.21.536599

**Authors:** YiRang Shin, Matthew R. Lowerison, Yike Wang, Xi Chen, Qi You, Zhijie Dong, Mark A. Anastasio, Pengfei Song

## Abstract

Ultrasound localization microscopy (ULM) is an emerging super-resolution imaging technique for deep tissue microvascular imaging. However, conventional localization methods are constrained by low microbubble (MB) concentration, as accurate localization requires a strict separation of MB point spread functions (PSFs). Furthermore, deep learning-based localization techniques are often limited in their ability to generalize to *in vivo* ultrasound data due to challenges in accurately modeling highly variable MB PSF distributions and ultrasound imaging conditions. To address these limitations, we propose a novel deep learning-pipeline, LOcalization with Context Awareness (LOCA)-ULM, which employs simulation that incorporates MB context to generate synthetic data that closely resemble real MB signals, and a loss function that considers both MB count and localization loss. In *in silico* experiments, LOCA-ULM outperformed conventional localization with superior MB detection accuracy (94.0% vs. 74.9%) and a significantly lower MB missing rate (13.2% vs 74.8%). *In vivo*, LOCA-ULM achieved up to three-fold increase in MB localization efficiency and a × 9.5 faster vessel saturation rate than conventional ULM.

## Main

Super-resolution optical microscopy is an established optical imaging technology that breaks the diffraction barrier of light and offers an order-of-magnitude improvement in imaging spatial resolution. One of the successful implementations of optical super-resolution, called single-molecule localization microscopy (SMLM), uses the stochastic blinking of fluorophore emissions to avoid overlaps between PSFs of individual molecules within a dense sample ^1, 2^. The sample is repeatedly illuminated, and a super-resolved image is reconstructed by accumulating the localized positions of single emitters that were separated in time. SMLM provides a nano-scale spatial resolution, which is essential for biological research at the cellular and subcellular levels ^3^.

The concept of localization microscopy has been successfully adopted by the ultrasound community to overcome the acoustic diffraction limit. As an acoustic analog to SMLM, ultrasound localization microscopy (ULM) uses ultrasound contrast agents (i.e., microbubbles or MBs) that flow within the blood vessels as individual point targets to achieve super-resolution ^4, 5^. By localizing each MB, ULM increases the ultrasound imaging spatial resolution by an approximate factor of ten ^6^. Because ULM uses the strong backscattering signal from MBs, it does not sacrifice imaging depth of penetration to gain spatial resolution. This key advantage makes ULM a powerful tool that allows noninvasive probing of deep tissue microvasculature for many preclinical and clinical applications ^7^.

As with all imaging techniques, ULM is not without limitations. At present, a key limitation of ULM is the long data acquisition time, which is the result of the inherent compromise between MB concentration and MB localization efficiency/accuracy. For example, to achieve accurate MB localization, ULM requires limited number of MBs per imaging frame (e.g., by using low MB concentration) so that MBs are spatially separated and localizable. However, lower MB concentration also makes it slower to accumulate adequate MB localizations to fill the lumen of the vessel, which can take several to tens of minutes ^8^. In contrast, a higher MB concentration speeds up the MB filling process in theory, but in practice it also increases MB signal overlap and reduces MB localization efficiency. As a result, increased MB concentration does not necessarily translate to faster ULM imaging speed. As such, improving MB localization efficiency under high MB concentrations remains a challenging yet essential task for improving the imaging speed (i.e., temporal resolution) of ULM.

Various methods have been proposed to improve MB localization under high-density MB conditions. Earlier studies focused on using Fourier-based filters to separate overlapping MBs into subgroups, leveraging the diverse spatiotemporal flow characteristics of MBs ^9^. Assuming a sparse distribution of MBs in each imaging frame, localization algorithms based on sparse image recovery and compressed sensing (CS) have been proposed to separate overlapping MB signals ^10–12^. However, the effectiveness of these methods depends on constructing an accurate MB PSF forward model, which is challenging due to the nonlinear response of MBs as well as other complexities involving frequency-dependent ultrasound attenuation, phase aberration, multi-scattering, and multi-path reverberation. In addition, the assumption of sparsity does not necessarily hold in areas with high MB density, which hampers the performance of these methods.

Deep learning has emerged as a promising solution for robustly localizing high-density MBs in ULM. One major limitation of deep learning-based MB localization is the lack of ground truth for the MB PSF in *in vivo* imaging settings. As a result, different MB PSF modeling techniques (e.g., using bivariate Gaussian models ^13^ and Field II simulations ^14^) have been proposed and commonly used to generate simulation datasets to train deep networks for MB localization. However, the complex MB acoustic responses *in vivo* ^15–17^ make it difficult to generate MB PSFs that closely resemble real MB signal characteristics (e.g., size, shape, brightness). Since the performance of deep learning localization heavily depends on the dataset that it is trained on, inaccurate estimation of the MB PSF can introduce biases in the model. Furthermore, existing simulation pipelines do not incorporate the appropriate MB signal properties and ultrasound system characteristics that are frequently observed from *in vivo* ultrasound data. As a result, the performance of existing deep learning-based ULM techniques is greatly undermined by the inaccurate modeling of the MB PSF and ultrasound imaging system.

In this work, we present LOcalization with Context Awareness (LOCA)-ULM, which leverages deep learning and contextual information to achieve robust localization under high MB concentrations. We first address the challenge of creating realistic synthetic datasets by constructing an MB PSF model informed by real ultrasound data. Considering the need for a flexible PSF simulator that can describe the high variability of MB PSFs, we utilize a generative adversarial network (GAN) ^18^ to directly learn the underlying properties of real *in vivo* MB signals. GANs are powerful tools for generating realistic samples without requiring domain-specific knowledge to model real data distributions. As a result, they are well-suited for capturing the complex properties of MB signals and creating realistic MB PSFs. Moreover, our simulation includes modeling of both ultrasound system noise and MB behavior. To accurately represent MB behavior, we assigned parameters related to factors such as brightness, lifetime, and velocity, to create ground truth positions. Overall, by simulating GAN-based MB signals that also reflect the *in vivo* MB behavior, we can better train the network to identify MB signals present in ultrasound images, leading to improved performance. The second method aims to address the challenges associated with high MB concentration that are present in practice. We translated the Deep Context Dependent (DECODE) ^19^ neural network into the ULM framework, in which the DECODE architecture and loss functions were optimized to achieve robust MB localization across a wide range of MB concentrations. In this paper, we systematically evaluated the performance of the proposed methods with both simulation and *in vivo* data that include chicken embryo chorioallantoic membranes (CAMs) and rat brains.

## Results

Fig. 1a illustrates the simulation pipeline designed to simulate realistic MB response and ultrasound imaging conditions. The simulation used MB PSFs generated by a Least-squares Generative Adversarial Network (LSGAN) ^20^. Conventional localization techniques have limitations in creating a complete set of MB PSFs that accurately represent the distribution of real MB PSFs. To overcome this challenge, we employed LSGAN (as depicted by “*G*” in Fig. 1a) to create a more extensive collection of MB templates that could be used for training the LOCA-ULM, including those that have not been observed during training. This strategy enables us to construct a more robust localization network that can detect various MBs with different shapes, leading to a enhance localization performance on *in vivo* ultrasound data at inference stage.

**Fig 1.**
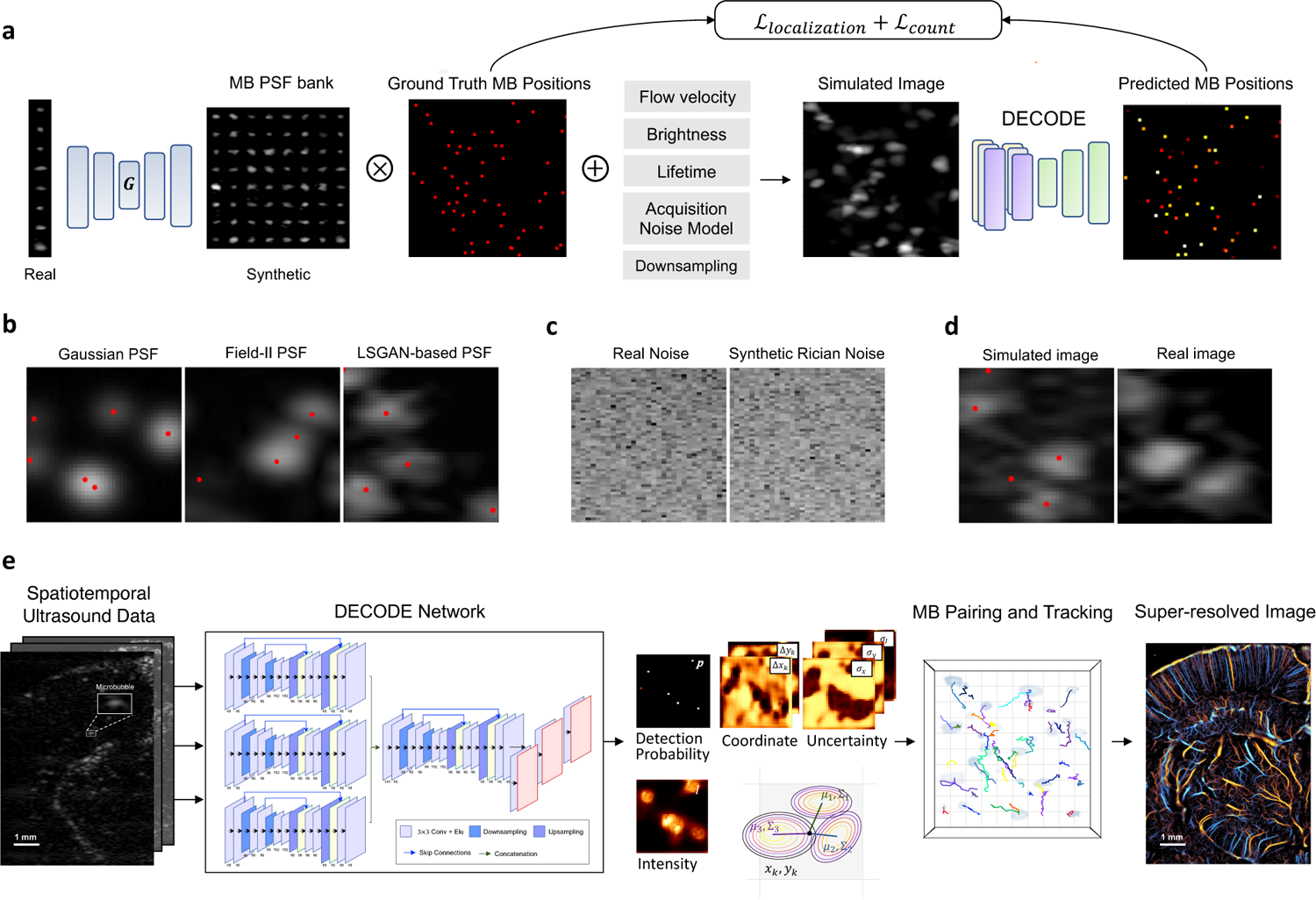
Overview of the proposed LOCA-ULM MB localization pipeline. LOCA-ULM is a simulation-based supervised learning method using MB PSFs generated by Least-squares Generative Adversarial Network (LSGAN)^20^. and DECODE localization^19^ **a** The LSGAN (*G*) was trained on a large set of MB PSFs identified by the conventional normalized cross-correlation (NCC) localization algorithm^21^. The LSGAN learns the distribution of real MB signals and generates diverse and realistic synthetic MB PSFs. The LSGAN-generated MB PSFs are convolved with simulated ground truth MB positions assigned with MB-specific characteristics (e.g., brightness, velocity, lifetime) to create simulated images that closely resemble real data. The simulated images were used to train the DECODE network for localization purposes. **b** Examples of simulated MB signals using different PSF simulation methods (2D Gaussian, Field II, and LSGAN). Red dots indicate the ground truth MB location. **c** Examples of experimentally acquired electrical noise from the ultrasound system, synthesized Rician noise using the proposed method (Methods). **d** Simulated image with LSGAN-based MB PSFs with added Rician noise and real MB image extracted from the *in vivo* CAM dataset. **e** DECODE-based ultrasound localization microscopy pipeline. Inference was performed by using *in vivo* ultrasound data. 2D-DECODE outputs the probability of detecting a MB near pixel *k* (p_k_), sub-wavelength spatial coordinates (Δx_k_, Δy_k_) respect to center of the pixel k, MB brightness (*I*), and corresponding uncertainties (σ_x_, σ_y_, σ_I_). MB pairing and tracking were applied to predicted coordinates and the final super-resolved ULM images were generated.

LSGAN was initially trained on a large number of MB signals obtained from *in vivo* images using a conventional localization algorithm based on normalized cross-correlation (NCC) ^21^. Once trained, the LSGANs were used to generate realistic MB PSFs that were stored in an MB PSF bank (i.e., collection of MB templates that were later used by DECODE network for training). To create the ground truth MB positions, a list of sub-wavelength coordinates was randomly sampled and assigned factors such as MB brightness, lifetime, and velocity to emulate real MB behavior. The ground truth positions were then convolved with the synthesized MB PSFs (randomly chosen from the MB PSF bank) to create ultrasound images with realistic MB signals. A typical simulated image based on LSGAN-created MB signal is shown in Fig. 1b (LSGAN-based PSF), which closely resembles real MB signals extracted from the *in vivo* CAM image shown in Fig. 1d (Real image), as compared to other MB simulation methods such as bivariate Gaussian and Field II. Finally, data-informed ultrasound noise was modeled and added to the simulated image to create the final training datasets for the DECODE network (Fig. 1c, d) (Methods).

In this study, we adopted the principles of DECODE to solve the problem of localizing spatially overlapping MB signals. We translated the cost functions of DECODE, including emitter count loss and localization loss, to train the network for the tasks of estimating MB counts, MB detection probability, MB locations, and MB brightness ^19^. The count loss and detection probability were jointly optimized with the localization loss to provide more accurate estimation of MB locations. This is a more robust approach than directly using MB location as loss terms (e.g., 1’s for the center of MBs and 0 for otherwise), which can generate a challenging condition for training ^13, 22^. We trained the network using simulated images using LSGAN-generated MBs, and the count loss and localization loss were optimized to maximize the likelihood of the ground truth MB positions under a Gaussian mixture model (LOCA-ULM). In the inference stage (Fig. 1e), the network estimates the true distribution of MB locations and brightness from real ultrasound data used as input. This strategy allows the network to output confident MB detection probability and accurate localizations for spatially overlapping MB PSFs (Methods).

### Simulation study

We first validated the proposed LOCA-ULM localization pipeline using simulation data. Using the simulation pipeline described in Methods, the test dataset was created using the MB signals extracted from *in vivo* CAM data. Two-thousand imaging frames with an image size of 100 pixels × 100 pixels (12.3 μm pixel size) were generated for different MB concentrations. The concentration for the simulation was incremented with a 0.02 MBs/λ^2^ step size from 0.02 MBs/λ^2^ (low density) to 0.37 MBs/λ^2^ (high density), where λ is the wavelength of the ultrasound pulse used for imaging (Table 1). Fig. 2a shows examples of MB localization using LOCA-ULM and conventional localization on simulation datasets with different MB concentrations. At a low MB concentration (0.06 MBs/λ^2^), both LOCA-ULM and conventional MB localization methods provided MB locations that agree with the ground truth. As the concentration increased (0.27 MBs/λ^2^), conventional localization started to miss more MBs (blue arrows in Fig 2a), and the miss-localizations tend to occur around the center of the clustered MB signals (yellow arrows in Fig. 2a). In contrast, LOCA-ULM was able accurately separate and localize overlapping MB signals with various shapes and brightness. LOCA-ULM localization is also in good agreement with the ground truth.

**Fig 2.**
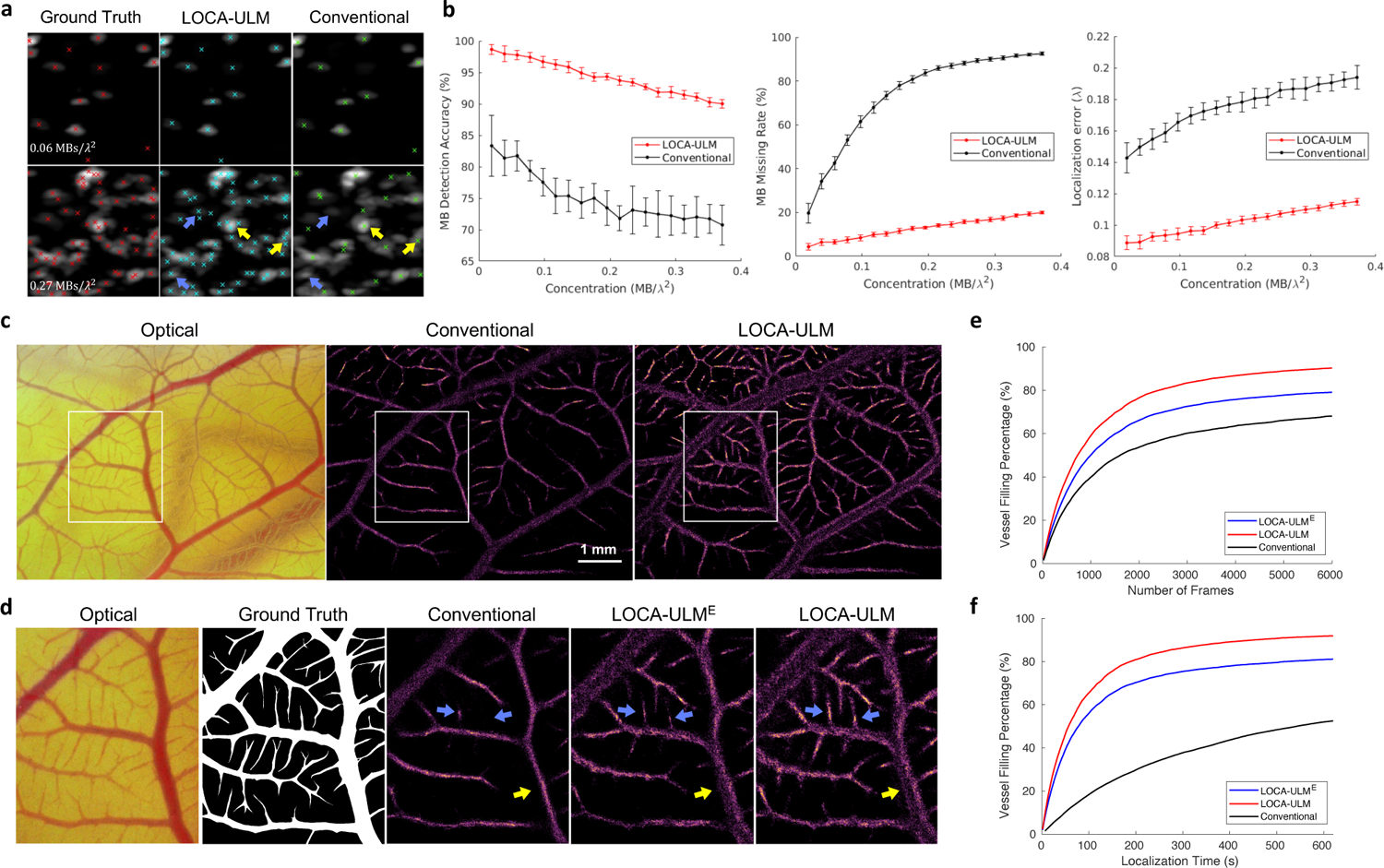
Results of the simulation study and in vivo chicken embryo CAM imaging study. **a** Simulation results of LOCA-ULM and conventional localization with low (0.06 *MBs*/λ^2^) and high (0.27 *MBs*/λ^2^) MB concentrations. Ground truth MB positions are marked by red ×, LOCA-ULM localization by cyan ×, and conventional localization by green ×. **b** Comparison between LOCA-ULM and conventional localization was performed on simulated images at increasing MB concentrations, using three performance metrics: MB detection accuracy, MB missing rate, and localization error (Methods). **c-f** Comparisons among conventional localization, LOCA-ULM^E^, and LOCA-ULM MB localization in *in vivo* chicken embryo CAM imaging. **c** Optical microscopy image of the CAM surface microvessel, along with MB localization images reconstructed by conventional localization and LOCA-ULM. **d** The ROI selected from the optical image and the corresponding ground truth vessel segmentation. Magnified view of the MB localization images marked by the white ROI for conventional localization, LOCA-ULM^E^, and LOCA-ULM. **e,f** The vessel filling (VF) percentage of three conventional localization, LOCA-ULM^E^, and LOCA-ULM, as a function of the number of frames and localization time.

### LOCA-ULM achieves high MB localization accuracy and efficiency under high MB concentrations in simulation data

The MB localization performance on simulation data was evaluated quantitatively using three metrics (Methods). Fig. 2b shows the performance of LOCA-ULM versus conventional localization with respect to increasing MB concentrations. LOCA-ULM consistently outperformed the conventional localization algorithm in regard to MB detection accuracy and MB missing rate, especially for high concentration conditions. LOCA-ULM achieved an average detection accuracy of 94.0%, as compared to an average accuracy of 74.9% from conventional localization. LOCA-ULM also substantially decreased the missing rate (13.2% vs 74.8%) over conventional localization. This improvement is essential for shortening the data acquisition time for ULM because it allows higher concentration MBs to be administered *in vivo* while maintaining a robust MB localization performance with high efficacy.

The localization error in Fig. 2b shows that LOCA-ULM consistently reduced the MB localization error across all concentrations when comparing to conventional localization. The theoretical resolution limit of ULM (i.e., localization error) can be estimated using the Cramér-Rao lower bound (CRLB)^23^. Following the theoretical CRLB model, we predicted a maximum resolution of 3.29 μm with the CAM study acquisition settings. In low-density conditions (0.02 MBs/λ^2^), conventional localization achieved a maximum localization resolution of 10.87 μm, while LOCA-ULM achieved localization resolution of 6.74 μm, which is closer to the CRLB.

### GAN-generated MB signals improved LOCA-ULM performance for MB localization in the in vivo CAM imaging study

The performance of LOCA-ULM was further evaluated in an *in vivo* CAM microvessel model. To demonstrate the effectiveness of using LSGAN-generated realistic MBs, we used two types of simulation data for network training. The first type (LOCA-ULM^E^; LOCA-ULM Experimental) used MB signals directly extracted from the CAM data; the second type (LOCA-ULM) used MB signals generated by the LSGAN. Fig. 2e summarizes the vessel filling (VF) percentage for all the localization methods including conventional localization, LOCA-ULM^E^, and LOCA-ULM (Methods). LOCA-ULM^E^ and LOCA-ULM achieved consistently higher VF percentage and faster vessel saturation rate than conventional localization. At 6000 frames (total 6 seconds of acquisition) for MB accumulation, LOCA-ULM achieved the highest VF percentage (90.25%), followed by LOCA-ULM^E^ (79.15%) and conventional localization (69.09%). Notably, the VF percentage respect to optical image ground truth of conventional localization started to plateau around 70%, while LOCA-ULM did not plateau until 90%. This result is consistent with the observation of under-filling of the major vessels using conventional localization as indicated by the yellow arrows in Fig. 2d. Both LOCA-ULM and LOCA-ULM^E^ filled the large vessels more completely and the size of the vessel was closer to the ground truth (i.e., based on optical microscopy). In Fig. 2d the reconstructed microvessels indicated by the blue arrows display higher intensity for LOCA-ULM, revealing vessel structures that have not yet been fully reconstructed by conventional localization and LOCA-ULM^E^. These results suggest that the diverse MB signals generated by the LSGAN enhanced the network’s capability of localization more MBs under high MB concentrations.

### LOCA-ULM significantly improves computational performance of MB localization

In addition to faster and more robust vessel filling performance, LOCA-ULM also enjoys an accelerated processing time over conventional localization, thanks to the high inference speed of deep neural networks. The LOCA-ULM network took 78s to localize 1000 ultrasound imaging frames with the size of 720 pixel× 560 pixel (7.10 mm × 5.52 mm), representing a 4-fold speedup over conventional localization. To achieve a 50% VF percentage, LOCA-ULM needed the least amount of ultrasound images (740 frames), which translates to the fastest localization time (57.81 s) over LOCA-ULM^E^ (1000 frames, 78.13s) and conventional (1620 frames, 546.75s) (Fig. 2f). These results indicate that LOCA-ULM greatly enhances the ULM performance by reducing both the data acquisition time (i.e., shorter MB accumulation) and post-processing time while providing higher MB localization efficacy.

### LOCA-ULM demonstrates superior in vivo ULM imaging performance in a rat brain

We demonstrated the generalizability of LOCA-ULM using *in vivo* rat imaging datasets. Fig 3. c-d shows the final ULM images based on 20000 frames (a total of 20 seconds of data acquisition) of accumulation. After localization, MB pairing and tracking were performed using uTrack^24^. As shown in Fig. 3d, the vascular bed in the rat brain presents large variations of vessel sizes with wide distribution of MB concentrations. As shown in Fig. 3c, conventional localization suffered from poor localization performance in regions with high MB concentrations, which manifest as disconnected and missing vessels in these regions (red arrows in Fig. 3e). In contrast, LOCA-ULM revealed the dense cerebral vascular networks in these regions, which were well-perfused and fully connected (red arrows in Fig. 3f).

**Fig 3.**
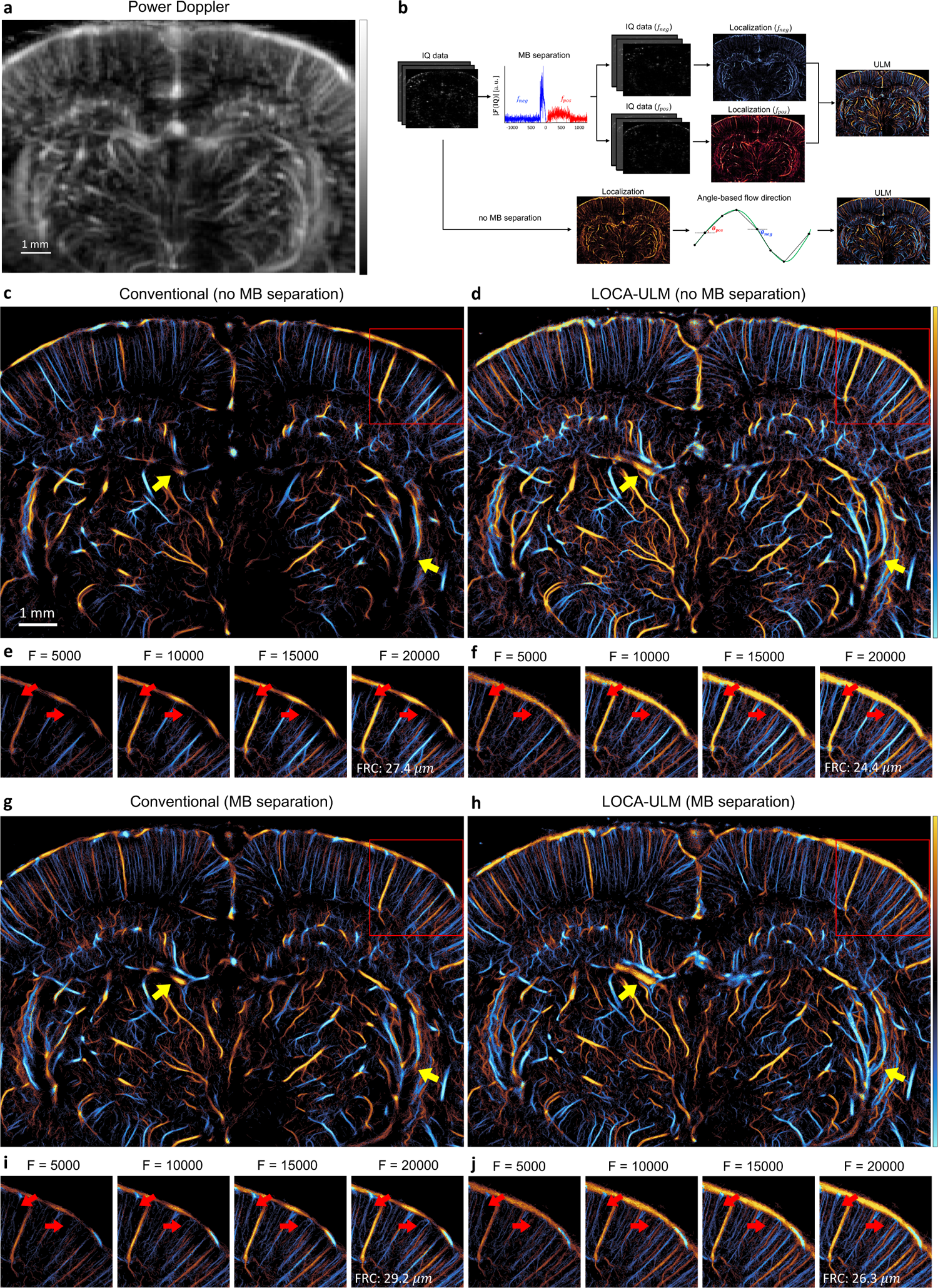
Comparison of LOCA-ULM and conventional localization to *in vivo* rat brain ultrasound data. **a** Power Doppler image generated by accumulating 2500 frames (a total of 2.5 seconds of acquisition) of rat brain ultrasound data. **b** *in vivo* rat brain localization workflow. The IQ data after tissue clutter filtering was processed with and without Fourier-based MB separation. For MB separation, the high concentration MB dataset was divided into subsets of upward and downward flow towards the transducer using a directional filter ^9^. Angle-based flow direction was used for dataset without MB separation. For each dataset, MB locations was determined by performing normalized cross-correlation with an empirically determined PSF function (i.e., conventional localization) or LOCA-ULM. The uTrack algorithm was used to pair the localized MB centers and estimate their trajectories. **c-j** Each ULM directional flow maps were generated by accumulating 20000 frames (a total of 20 seconds of acquisition), **c,d** without MB separation and **g,h** with MB separation. **e,f,i,j** Improvement of vessel structures respect to the increasing number of frames is displayed on the bottom, shown for the area marked with red rectangle. F indicates number of frames used for ULM reconstruction and FRC indicates Fourier Ring Correlation.

Next, we designed a study to compare LOCA-ULM with the *state-of-the-art* MB localization method based on MB separation ^9^. We used the MB separation filter to separate the ultrasound MB data into two subgroups: flow away from the transducer (downward flow) and flow toward the transducer (upward flow), as shown in Fig. 3b. Fig. 3g, h demonstrate that MB separation facilitated more robust MB localization and tracking in high MB density regions for both conventional and LOCA-ULM. The improvement is most significant for conventional localization, which suffered from poor MB localization performance in high density MB regions. The intersecting and adjacent small vessels that were missing by conventional localization now become clearly visible by using MB separation. For LOCA-ULM, the improvement was moderate because LOCA-ULM was already efficient with localizing MBs in high density regions. This is evidenced by comparing Fig. 3d and Fig. 3h where most of the cerebral vasculature was consistent before and after applying MB separation for LOCA-ULM (indicated by yellow arrows). When comparing Fig. 3d and 3g, it becomes clear that even with MB separation, conventional localization still could not achieve a similar MB localization performance to LOCA-ULM without MB separation. This is an important finding because it suggests that LOCA-ULM alone can already outperform the state-of-the-art MB localization technique and does not require the help of post-processing methods such MB separation.

Finally, the spatial resolution of the ULM reconstructions were characterized by applying the Fourier Ring Correlation (FRC) method, using the track splitting strategy and a 2-σ threshold curve as proposed by Hingot et al ^25^. Our results showed that both LOCA-ULM and conventional localization produced spatial resolution below a half wavelength at the imaging frequency 15.625 MHz (49.28μm) with and without MB separation (Supplementary Fig. 1). These findings suggest that LOCA-ULM can achieve a more complete reconstruction of the vascular network and provide visualization of well-perfused vessels without compromising spatial resolution.

### LOCA-ULM-based MB localization automatically adapts to different MB concentrations

To further investigate the performance of LOCA-ULM under varying MB concentrations *in vivo*, we conducted an experiment with increasing MB injection rate from 20 μL/min to 40 μL/min (Methods). Fig. 4 shows the reconstructed ULM images for 20 μL/min and 40 μL/min injection rate using conventional and LOCA-ULM localization in a rat brain, where a total of 25000 frames (a total of 25 seconds of acquisition) of ultrasound data were used for reconstruction. Similar to the observations in Fig. 3, LOCA-ULM demonstrates much more complete cerebral vasculature reconstruction than conventional localization. Notably, vessel areas with high MB concentrations suffered from the high MB missing rate of conventional localization, which disappeared in the ULM image (red arrows in Fig. 4a, c). In contrast, LOCA-ULM revealed large vessel structures that were missed by conventional localization (red arrows in Fig 4b, d). Furthermore, in regions with moderate MB concentration, the intensity of reconstructed vessels with conventional localization decreased with an increase in MB injection rate, leading to degraded vessel delineation. For example, for conventional ULM, the two close-by vessels that were separable at an MB injection rate of 20 μL/min (yellow arrows in Fig. 4a) became indistinguishable at an MB injection rate of 40 μL/min (yellow arrows in Fig. 4c). LOCA-ULM significantly improved the ability to resolve adjacent structures, producing a clear separation of the vessels in both low (20 μL/min) and high (40 μL/min) MB injection rates (yellow arrows in Fig. 4b, d, respectively). LOCA-ULM also revealed tiny vessels near the cortical surface that cannot be reconstructed by conventional ULM, as indicated by green arrows in Fig. 4a-c.

**Fig 4.**
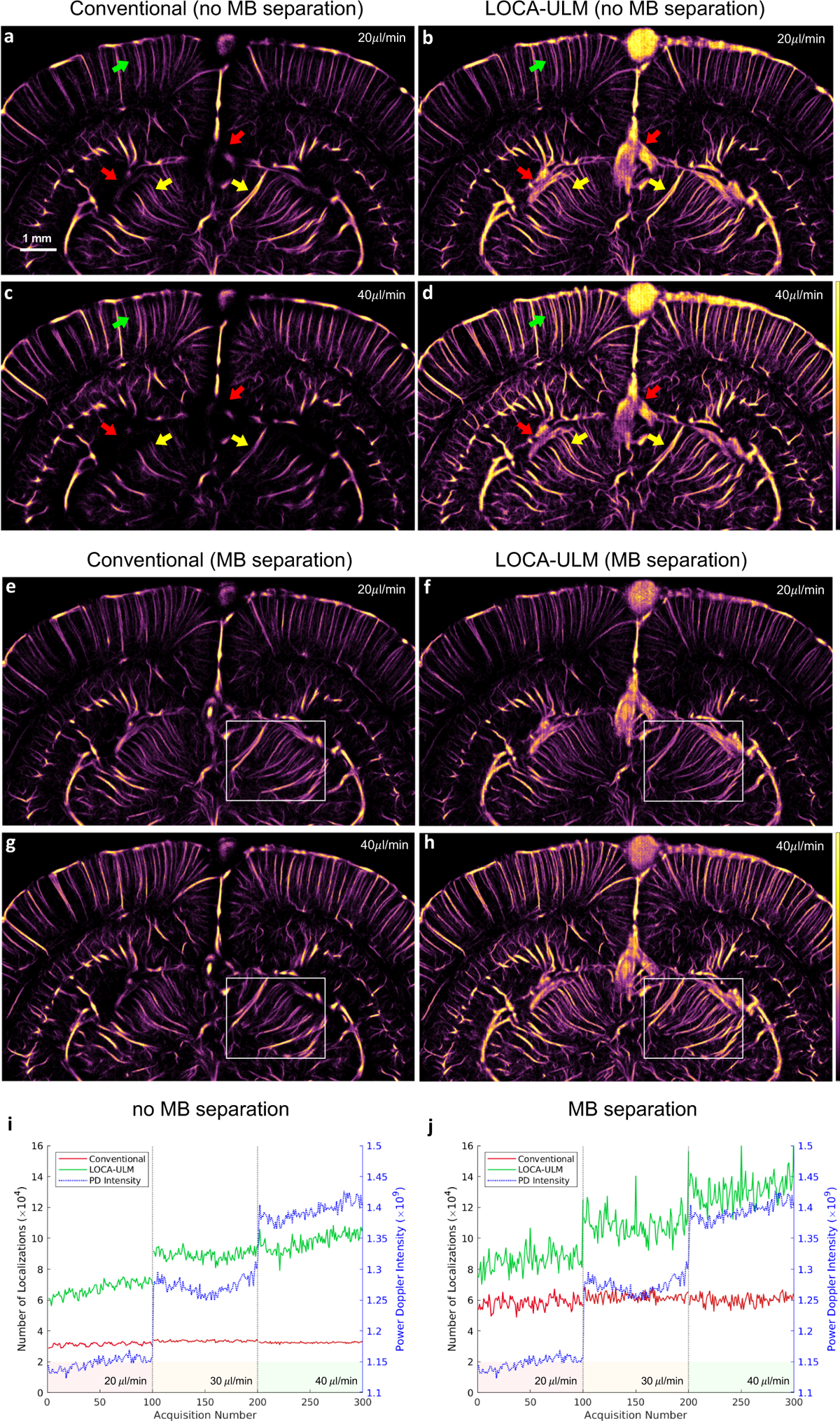
Effect of different MB injection rates (20 μL/*min*, 30 μL/*min*, 40 μL/*min*) on LOCA-ULM and conventional localization for rat brain ULM imaging. **a-h** Each ULM image was generated by accumulating 25000 frames of ultrasound data (a total of 25 seconds of acquisition) for MB injection rate of 20 μL/*min* and 40 μL/*min*, **a-d** ULM reconstruction without MB separation and **e-h,** with MB separation. **i,j** Comparison of total MB count per acquisition (a total of 250 frames per acquisition) for LOCA-ULM and conventional localization at different MB injection rates (20 μL/*min*, 30 μL/*min*, and 40 μL/*min*). Two datasets, **i** without MB separation and **j** with MB separation.

The performance of LOCA-ULM localization was further evaluated by a quantitative analysis that used the mean Power Doppler (PD) intensity as the reference. As shown in Fig. 4i, the MB count of LOCA-ULM closely followed the trend of increasing PD intensity while conventional localization did not. This result clearly indicates that conventional localization has already become saturated even at the lowest MB injection rate (20 μL/min). With the addition of MB separation, as shown in Fig. 4j, conventional localization improved the localization efficacy but was still saturated at the lowest injection rate. For LOCA-ULM, MB separation also improved the MB count, which suggests that there was missed localization for LOCA-ULM. Nevertheless, the improved MB count did not result in significant ULM image quality improvement (e.g., Fig. 4d, h). Finally, the quantitative results provide a good agreement with the ULM images, where LOCA-ULM reconstructed ULM images show increased microvessel intensity with increased MB injection rate, while conventional localization shows constant microvessel intensity despite the increased MB injection rate (white ROIs in Fig. 4 e-h).

## Discussion

In this study, we have presented LOCA-ULM, a context-aware deep learning-based MB localization method along with an LSGAN-based MB simulation pipeline to facilitate high quality ULM imaging under high MB concentrations. We adopted the principles of DECODE and designed a new simulation workflow that incorporated MB characteristics and realistic ultrasound imaging noise. Compared with well-established image formation processes of SMLM (e.g., blinking fluorophores, PSF modeling, noise and camera model) ^26^, there are several key differences to note in this study. First, the high variability of ultrasound MB PSFs makes it challenging to construct effective PSF models that fully and accurately capture the complexity of real MB signals. When trained with simple Gaussian PSF models, we immediately observed suboptimal ULM imaging performance (e.g., gridding artifacts, poor vessel reconstructions) (Supplementary Fig. 2). The proposed LSGAN-based MB generation implicitly learns the complex MB PSF distributions that are present in the given ultrasound data, effectively minimizing the discrepancy between real and modeled MB signal. We demonstrated the advantages of LSGAN-generated PSFs in *in vivo* CAM imaging, where LOCA-ULM demonstrated significantly higher vessel filling (VF) percentage and faster vessel saturation rate over LOCA-ULM^E^ and conventional localization (Fig. 2e, f). Secondly, we demonstrated the adaptiveness and robustness of LOCA-ULM with context-aware training for varying MB concentrations. Unlike SMLM, where emitter density can be reduced using laser irradiation or by adjusting chemical environment ^26^, controlling MB concentration for ULM is challenging due to the wide distribution of vessel sizes and hemodynamics *in vivo*. As a result, the high missing rate and inaccuracy of conventional localization leads to incomplete ULM reconstructions with corrupted and missing vessel structures (Fig. 3c, Fig. 4a, c). Deep learning offers a practical solution for learning the non-linear mapping from ultrasound images with overlapping MB signals to sparse localization maps in a data-driven manner. However, the sparse nature of localization maps hinders the direct training of deep networks using common loss functions based on least-squares regression under the *l*_1_ regularization ^13, 27^. In our study, we leveraged the joint optimization of both MB count loss and localization loss into the training process to achieve accurate MB localization in both low and high MB densities. The count loss encourages the network to output a sparse and high probability detection map, providing a complementary information about the position of each MBs. In turn, the localization loss models the localized MB centers as the sum of Gaussian distributions weighted by the detection probability. It was shown in the simulation study that LOCA-ULM is highly capable of separating overlapping MB signals, resulting in the best match with the ground truth positions at high MB concentration (Fig. 2a).

Furthermore, to provide the network with additional context of the MB signals, we integrated MB-specific characteristics such as brightness, movement, lifetime, and ultrasound noise to our simulation framework (Fig. 1a). This enables LOCA-ULM to comprehend the distinct features of real MB signals, which enhanced the localization precision and ULM image quality. Our simulation study showed that LOCA-ULM yielded superior MB localization efficiency compared to conventional localization, improving detection accuracy (94.0% vs. 74.9%) and reducing the missing rate (13.2% vs 74.8%). In our *in vivo* CAM results, LOCA-ULM achieved a more complete filling of larger vessels and reconstructed microvessels with higher intensity, as validated by optical imaging (Fig. 2d). In *in vivo* rat brain study, LOCA-ULM was able to maintain high localization accuracy even with considerable increase in MB injection rate (i.e., 40 μL/min), with up to a three-fold increase in detected MB localizations compared to conventional localization (Fig. 4i). Likewise, LOCA-ULM reconstructed well-connected and perfused cerebral vasculature, including large and densely populated vessels missed by the conventional ULM (Fig, 3d, Fig. 4b, d). LOCA-ULM also reconstructed more vessels with higher fidelity at high MB concentration, especially the adjacent microvessels that could not be resolved using conventional localization (Fig. 4b, d).

We have also demonstrated the effectiveness of LOCA-ULM in achieving both high-speed processing and accelerated data acquisition for ULM. In theory, higher MB concentration is necessary for faster ULM imaging because it accelerates vessel filling rate of smaller vessels of MBs, which translates to shorter data acquisition time ^8^. Our results indicate that LOCA-ULM achieved increased MB count in line with increased MB injection rate (Fig. 4i, j), enabling accelerated acquisition with faster MB perfusion. Moreover, we demonstrated that the quality of the reconstructed ULM images for LOCA-ULM was not significantly affected by MB separation (Fig. 3d, h and Fig. 4 d, h). This result is significant because it indicates that LOCA-ULM can substantially reduce the ULM post-processing time by eliminating the need of dividing datasets into subsets with sparser MB concentration.

This study has another notable aspect, which is that LOCA-ULM can be easily applied to a broad range of ultrasound imaging scenarios. Our proposed simulation pipeline does not require any prior knowledge of the PSF model or the ultrasound image formation process to create the training dataset. This is not the case for most deep-learning based localizations, which typically necessitate specific knowledge of imaging factors, such as Field II simulation parameters ^14^ or 2D Gaussian PSF model ^27^, to generate simulated dataset. Our method can be easily used to create simulated data, which can aid in robust training and reduce the challenge of generalizing deep learning-based localization to *in vivo* ultrasound data.

Our study has some limitations. First, the DECODE network and LSGAN need to be retrained when the ultrasound imaging settings are altered. In addition, a stable training of LSGAN requires a large collection of spatially isolated MB signals extracted from experimental data. Also, the performance of LOCA-ULM may be undermined by inaccurate simulation parameters (e.g., MB brightness, background noise, etc.), resulting in suboptimal MB localization performance. Nevertheless, because LOCA-ULM outputs uncertainties of localizations, one can use the predicted uncertainties to reject unreliable localizations. Finally, the input ultrasound image to LOCA-ULM needs to be upsampled to avoid quantization artifacts. This results in an overall increased computational cost for the proposed localization technique.

## Methods

### Simulation Pipeline

The simulated datasets for training are generated during the network training, creating 10000 frames per epoch, and using each frame only once for training. Because LOCA-ULM is trained purely on simulated data, it may fail to generalize to real ultrasound data if there is a discrepancy between the two datasets. To address this issue, we created a realistic model for the ultrasound image formation process that incorporates PSF model based on LSGAN and data-informed ultrasound noise (Fig. 1a). Compared with the generic GAN, LSGAN replaces the sigmoid cross entropy loss in the discriminator with a least squares loss, facilitating the generator to create more realistic images and learn the distribution of the training data more robustly ^20^. LSGAN has been applied in medical imaging to improve spatial resolution and prevent mode collapse (i.e., generator creating limited ranges of outputs)^28–30^. The training problem for the LSGAN can be formulated as:

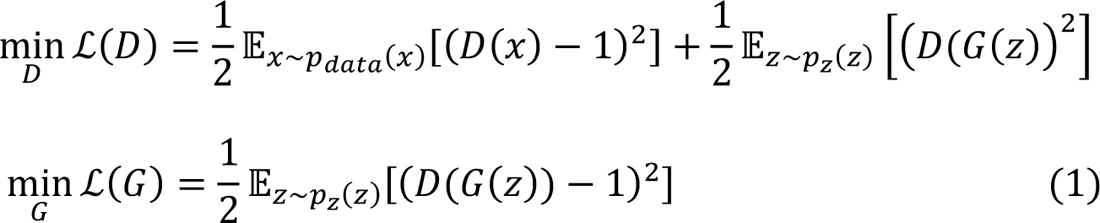

where *D* denotes the discriminator, *G* represents the generator, *Z* represents the input signal, which was randomly sampled from a uniform distribution, and x represents the MB PSFs extracted from real ultrasound images.

To collect the LSGAN training data, the in vivo ultrasound images were first interpolated by a factor of 5 (5X) in axial dimension and 10X in the lateral dimension. This corresponds to a 0.064 λ pixel size for the CAM images and 0.1 λ pixel size for rat brain images (Table I, PSF pixel resolution). Square patches (65 *pixel* × 65 *pixel*) were extracted from the in vivo ultrasound images and used to create the simulated images for training. Each patch contains a single MB PSF that takes the peak location identified by the normalized cross-correlation (NCC) localization algorithm as the true MB location ^21^. A total of 3000 patches were manually selected from the in vivo ultrasound images to train the LSGAN and the mean (μ_I*max*_) and standard deviation (σ_I*max*_) of the maximum intensity were calculated. After training, the synthetic PSFs generated from the LSGAN were saved into a bank of PSFs. To generate training data for the network, a list of ground truth MB positions was sampled in sub-wavelength pixel resolution (Table I, DECODE output pixel resolution) and convolved with randomly selected MB PSFs retrieved from the PSF bank. The brightness of MB PSFs was drawn from a Gaussian distribution *N*(μ_I*max*_, σ_I*max*_). To generate diverse simulated frames, the first appearance of each MB was randomly selected from a continuous distribution, and the lifetime of each MB is chosen at random. 80 *pixel* × 80 *pixel* sized simulated frames were created and the images are down-sampled by a factor of 2 to create the final 40 *pixel* × 40 *pixel* sized training frames (Fig. 2b, LSGAN-based PSF).

To add realistic electronic noise to the simulation, we used Rician distribution as the noise model in this study. Assuming an additive Gaussian noise in both real and imaginary parts of the in-phase quadrature (IQ) data, the B-mode signal *I_x,z_*_,)_ (i.e., magnitude of IQ at pixel (x, *Z*)) satisfies the distribution:

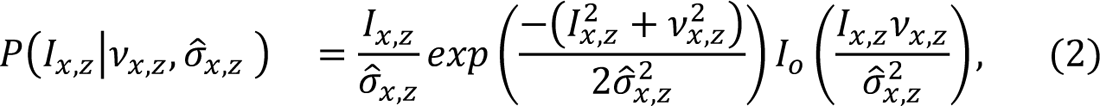

where ν*_x,z_*, is the magnitude of the B-mode signal at pixel (x, *Z)* without noise, σ*_x,z_*_)_ is the standard deviation of the additive noise, and *I*_o_ is the modified Bessel function of the first kind with order zero. In this study, the σ*_x,z_* was estimated experimentally by taking the temporal mean of the acquired electronic noise data *E(x, Z, t)* as,

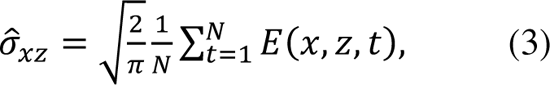

where *N* is the number of samples considered for estimation. Electronic noise in ultrasound images were obtained by performing the same ultrasound acquisition as the in vivo experiment without any imaging target (e.g., in air) (Fig. 1c).

### DECODE Architecture

Accurate and robust MB localization under a wide range of vessel sizes and MB concentrations is essential for successful ULM. Inspired by the previous study by Speiser, A. et al ^19^, we implemented the DECODE network that enables simultaneous detection and localization of MBs in a probabilistic framework. Several key aspects allow DECODE to outperform conventional localization methods. First, DECODE can improve detection and localization accuracy by capturing the temporal context of the MB flow. The architecture is divided into two networks: a *frame analysis network* that comprises three separate U-Nets, where features of three consecutive frames are extracted in each U-net. The frame analysis network is followed by a *temporal context network*, where the final outputs of the three U-Nets are combined to capture the temporal context information between neighboring frames (Fig. 1e).

Moreover, the DECODE network was trained to minimize the total loss that consists of three parts: an MB count loss (ℒ_count_), MB localization loss (ℒ_loc_) and a background loss (ℒ_bg_)^19^. The MB count loss is represented by a Bernoulli distribution p_k_ that indicates the probability of detecting a bubble near pixel k. Given that the probability p_k_ varies among the pixels, the mean (μ_count_) and variance (σ_count_^2^) of the Poisson-binomial distribution is given as μ_count_ = ∑_k_^K^ p_k_, σ_count_^2^ = ∑_k_^K^ p_k_ (1 – p_k_), where *N_GT_* is the total number of pixels. When *N_GT_* is sufficiently large, the Poisson binomial distribution approximates the Gaussian distribution defined as,

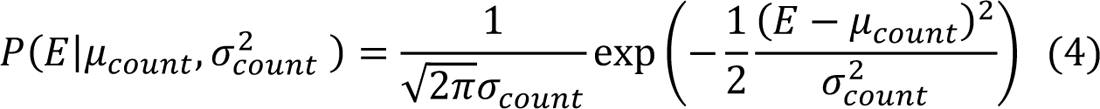

where E is the true number of simulated MBs. The log probability of E is maximized when the μ_count_ approximates to E, equivalent to minimizing,

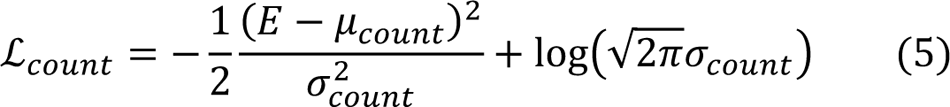

The localization loss is designed jointly to optimize the output variables of the Gaussian mixture model (GMM) to approximate to the true posterior with respect to MB locations and brightness. A GMM for each pixel k, weighted by the detection probability is used to approximate the true posterior. The four-dimensional Gaussian *P*(*u*_k_|μ_k_, Σ_k_) is modeled as a distribution over the coordinates and brightness of the MB *u* = [*x, y, Z, I*]:

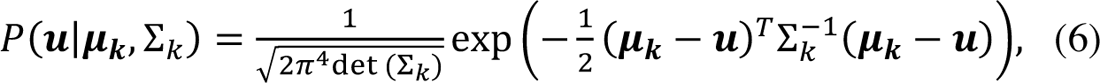

where μ_*k*_ = [x_k_ + Δx_k_, y_k_ + Δy_k_, *Z*_k_ + Δ*Z*_k_, *I*_k_] and Σ_k_ = diag(σ_x,k_^2^, σ_y,k_^2^, σ_z,k_^2^, σ_I,k_^2^). The (x_k_, y_k_, *Z*_k_) coordinate represents the center of pixel k, and (Δx_k_, Δy_k_, Δ*Z*_k_) is the sub-wavelength coordinates of the MB respect to the center of pixel k. The distance between the inferred posterior and the true posterior is minimized (i.e., by minimizing the forward KL divergence) by optimizing the log-likelihood of the weighted Gaussian distributions over the ground truth (GT) MBs,

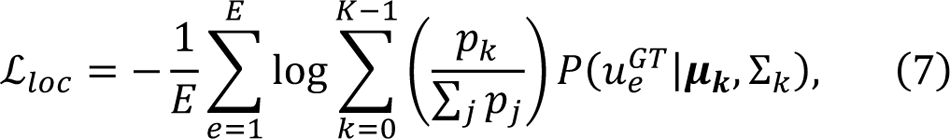

where e represents each MB present in the image. The localization loss maximizes the likelihood of the ground truth positions and brightness *u*_e_^GT^over all predicted detections. The DECODE network was designed to output the 9 parameters of the weighted Gaussian distribution respect to the center frame of the three consecutive imaging frames: (1) probability p_k_ that a MB was detected near pixel k, (2) the relative coordinates of the localized center Δx_k_, Δy_k_, Δ*Z*_k_ respect to the pixel center (x_k_, y_k_, *Z*_k_) (3) estimated brightness of the MB (*I*) (4) the uncertainties σ_x,k_, σ_y,k_, σ_z,k_, σ_I,k_, and (5) the background intensity (*B*). In this study, we used a 2D variant of DECODE to process the 2D ultrasound data. Also, the background loss (ℒ_bg_) in DECODE was removed since the background in ultrasound images was modeled separately using the noise model.

The DECODE network in Fig. 1e reveals the detailed architecture, where the U-Nets in the frame analysis and temporal context networks consist of two down-sampling and up-sampling layers. The convolution blocks in both networks adopted a kernel of 3 × 3 size followed by an Exponential Linear Unit (ELU) as an activation layer. The number of filters increases from 48, 96, and 192 for each down-sampling layer, with the feature map size halved. The number of filters decreases from 192, 96, and 48 for each up-sampling layer, with the feature map size doubled. The input of DECODE network were ultrasound images up-sampled to 2.5 X in axial dimension and 5 X in lateral dimension (Table I, DECODE network input pixel resolution).

### Evaluation Metrics

We compared three evaluation metrics to measure the MB localization performance of LOCA-ULM and conventional localization on simulation study. MB detection accuracy measures the fraction of correct localizations (within 5 pixels or 0.32λ the ground truth position) among all localized MBs:

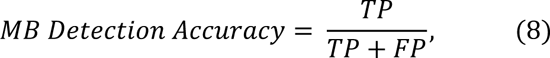

where TP is true positives and FP is false positives. The MB miss rate measures the fraction of missed localizations among all ground truth positions:

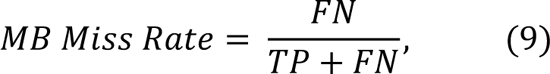

where FN is false negative. The localization error (*L*) computes the averaged root mean squared distance between the correctly localized MBs (i.e., TP) and the corresponding ground truth MB positions.

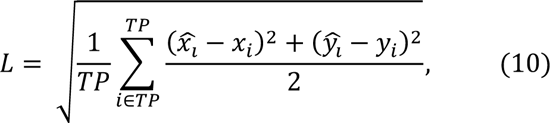

where x_i_, y_i_ are the ground truth coordinates and x_i_, yi are the predicted coordinates.

For quantitative assessment of the localization performance in *in vivo* CAM imaging, we calculated the vessel filling (VF) percentage using the method described by Kim, J. et al ^31^. First, a region of interest (ROI) that provided matching microvascular images between optical microscopy and ULM was selected. The optical image was resized with respect to the ULM image resolution to ensure accurate registration between the two images. The vessel structures in the optical microscopy image were carefully labeled by manual segmentation and used as the ground truth. The vessel filling (VF) percentage was calculated as

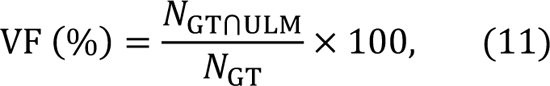

where *N*_GT_ is the total number of pixels classified as the ground truth vessels in the optical image. *N*_ST⋂VWX_ is the total number of pixels correctly classified by ULM with respect to the ground truth *N*_ST_.

### ULM Implementations

For each ULM data, an SVD-based clutter filter was applied to extract the MB signal from the surrounding tissue ^32, 33^. To reduce the intensity variations of the MB signal, all frames were normalized to a scale of 0 to 1 with respect to the minimum and maximum intensity within each acquisition (1600 frames for CAM study and 250 frames for rat brain study). Also, due to the hyperechogenicity of MBs, thresholding between the values of 0.1 − 0.2 was selected empirically to remove low-intensity background and noise. After image processing, the images were up-sampled to avoid the quantization artifacts associated with DECODE localization ^19^. Then, the network was trained to output super-resolved locations with sub-wavelength resolution (Table I, DECODE network output pixel resolution). For conventional ULM, normalized cross-correlation based MB localization was employed using a pre-defined multivariate Gaussian PSF ^21^. The centroid coordinates were input into the uTrack algorithm ^24^ for MB pairing and tracking, following a similar process as in our recent studies ^34^.

#### In vivo ULM data acquisition

1. Chicken Embryo Chorioallantoic Membrane (CAM) study We used the CAM microvessel model with optical imaging as ground truth to study the performance of different localization methods. Fertile chicken eggs were obtained by the University of Illinois Poultry Research Farm and kept in tilting incubators (Digital Sportsman Cabinet Incubator 1502, GQF Manufacturing Inc., Savannah, Georgia). After four days, the eggshells were removed, and the CAM embryos were mounted into a plastic holder in a position suitable for imaging. Then, the embryos were incubated for an additional 13 days in a humidified incubator (Darwin Chambers HH09-DA) until the desired developmental stage. A borosilicate glass tube (B120-69-10, Sutter Instruments, Novato, CA, USA) was pulled at high temperature and cut using a PC-100 glass puller (Narishige, Setagaya, Japan) to create a fine glass capillary needle for MB injection. 50 μL boluses of Definity^®^ solution (Lantheus, Bedford, MA) were injected into the surface bloodstream of the CAM via the glass needle. A high-frequency linear array transducer (L35-16vX, Verasonics Inc., Kirkland, WA) was placed at the side of the plastic holder to image the surface of the CAM vasculature. Ultrasound data were obtained by using a 9-angle compounding plane-wave imaging sequence (step size of 1°) with a center frequency of 20 MHz, pulse repetition frequency (PRF) of 40 kHz, and a post-compounding frame rate of 1,000 Hz. IQ data of 1600 frames per acquisition with a total of 20 acquisitions were generated (total 32 seconds of acquisition). Ground truth optical images were obtained using a Nikon SMZ800 stereomicroscope (Nikon, Tokyo, Japan) with A DS-Fi3 digital microscope camera (5.9-Mpixel CMOS image sensor, Nikon).
2. Rat Brain Study Ten-week-old Sprague Dawley rats (Charles River Laboratories, Inc.) were used in this study. Animals were anesthetized with isoflurane (5% induction, 1.5% maintenance) throughout the experiment. Before craniotomy, the jugular vein was catheterized and then the animal was fixed on a stereotaxic frame. The scalp was removed, and the skull was thinned using a rotary micromotor with a 0.5mm drill bit (Foredom K.1070, Bethel, CT). The skull was removed with the size of the cranial window of 12mm (left-right) by 6mm (rostral-caudal) below the bregma. To image the rat brain, Definity^®^ MBs were diluted with saline to yield an initial concentration of 1.44× 10^9^ bubbles per ml. The diluted MBs were continuously infused using a syringe pump (NE-300, New Era Pump Systems Inc., Farmingdale, NY). For the study of comparing the performance of different localization methods in different MB concentrations, we used an injection rate of 20, 30, 40 μL/min, and waited for 3 minutes after changing the injection rate to stabilize the systemic MB concentration. This corresponds to 1.8× 10^6^, 2.7× 10^6^, and 3.6× 10^6^ bubbles per ml of blood per minute, respectively. All rat brain data were acquired using a high-frequency linear array transducer (L22-14vX Verasonics Inc., Kirkland, WA). Ultrasound data were obtained by using a 5-angle compounding plane-wave imaging sequence (step size of 1°) with a center frequency of 15.625MHz, PRF of 28.57 kHz, and post-compounding frame rate of 1,000 Hz. IQ data of 250 frames per acquisition with total of 100 acquisitions were generated (total 25 seconds of acquisition). All procedures described above were approved by the Institutional Animal Care and Use Committee (IACUC) at the University of Illinois Urbana-Champaign. Details of the *in vivo* data acquisition specifications and image resolution are summarized in Table I.

## Author contributions

YS and PS designed and wrote the paper. YS and XC designed the simulation study. YW, QY, and PS prepared the rat model and performed craniotomies and ultrasound imaging on rat. MRL prepared the CAM model and performed ultrasound imaging. MRL designed the noise model. ZD designed the ultrasound transducer holder and programmed the motorized imaging stage. YS, MAA, and PS developed and applied the super-resolution ULM algorithm.

## Materials & Correspondence

Correspondence to Pengfei Song

## Data Availability

The data that support the findings of this study are available from the corresponding authors on request.

## Code Availability

The deep-learning models were developed based on DECODE available in Pytorch (https://github.com/TuragaLab/DECODE). Custom code for deployment of the simulation pipeline, LOCA-ULM training, and inference are available for research purposes from the corresponding author upon request.

## Supplementary Tables

**Table I.** *In vivo* study acquisition parameters and image resolution

|  | CAM study | Rat Brain Study |
| --- | --- | --- |
| **In vivo study acquisition parameters** |  |  |
| Transducer type | L35-16vX | L22-14vX |
| Center Frequency | 20 MHz | 15.625 MHz |
| Sampling Frequency | 125 MHz | 62.5 MHz |
| Wavelength | 77 $\mu m$ | 98.56 $\mu m$ |
| **Image resolution** |  |  |
| PSF pixel resolution | 4.928 $\mu m$ | 9.856 $\mu m$ |
| DECODE network input pixel resolution | 9.856 $\mu m$ | 19.712 $\mu m$ |
| DECODE network output pixel resolution | 4.928 $\mu m$ | 9.856 $\mu m$ |

