## Supplemental Figure 1,2 for "Context-Aware Deep Learning Enables High-Efficacy Localization of High Concentration Microbubbles for Super-Resolution Ultrasound Localization Microscopy"

### Supplementary Figures

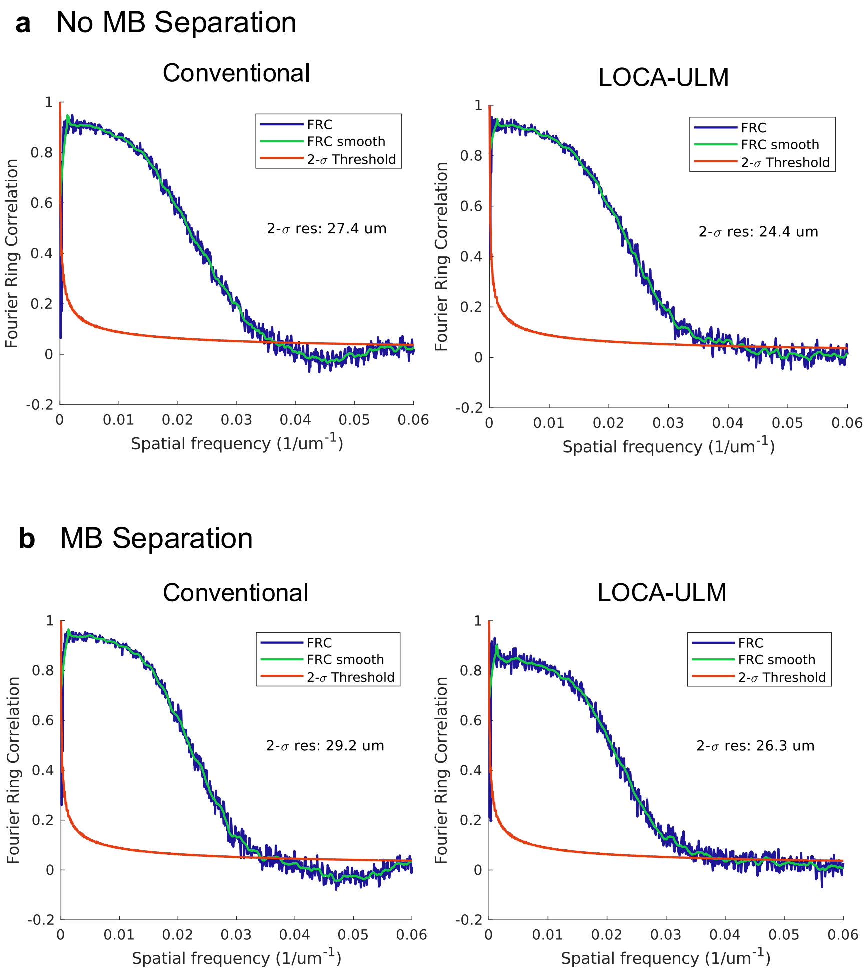


Fig 1. FRC curve using 2-$\sigma$ threshold for four ULM reconstruction. **a** Conventional and LOCA-ULM without MB separation. **b** Conventional and LOCA-ULM using MB separation.

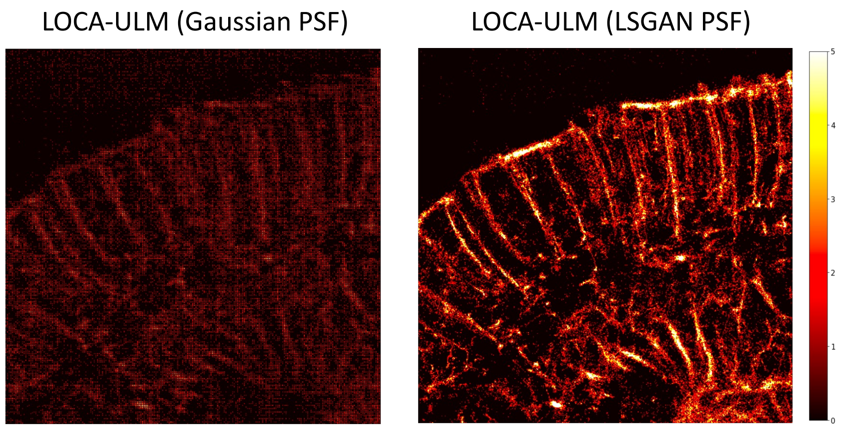


Fig 2. Comparison of LOCA-ULM localization results trained with different MB PSF model for simulation data. Gaussian PSFs and LSGAN-generated MB PSFs were used to create the simulated data for training. In vivo rat brain images were used for inference.
